# The apparent loss of PRC2 chromatin occupancy as an artefact of RNA depletion

**DOI:** 10.1101/2023.08.16.553488

**Authors:** Evan Healy, Qi Zhang, Emma H. Gail, Samuel C. Agius, Guizhi Sun, Michael Bullen, Varun Pandey, Partha Pratim Das, Jose M. Polo, Chen Davidovich

## Abstract

RNA has been implicated in the recruitment of chromatin modifiers, and previous studies have provided evidence in favour and against this idea. RNase treatment of chromatin is a prevalent tool for the study of RNA-mediated regulation of chromatin modifiers, but the limitations of this approach remain unclear. RNase A treatment during chromatin immunoprecipitation (RNase-ChIP or rChIP) reduces chromatin occupancy of the H3K27me3 methyltransferase PRC2. This led to suggestions of an “RNA bridge” between PRC2 and chromatin. Here we show that RNase A treatment during chromatin immunoprecipitation leads to the apparent loss of all facultative heterochromatin, including both PRC2 and H3K27me3 genome wide. This phenomenon persists in mouse embryonic stem cells, human cancer cells and human-induced pluripotent stem cells. We track this observation to a gain of DNA from non-targeted chromatin, sequenced at the expense of DNA from facultative heterochromatin, which reduces ChIP signals. Our results point to substantial limitations in using RNase A treatment for mapping RNA-dependent chromatin occupancy and invalidate conclusions that were previously established for PRC2 based on this assay.

**Highlights:**

- RNA degradation during ChIP-seq is insufficient to displace PRC2 from chromatin.
- RNA degradation led to the artificial depletion of ChIP-seq signals in multiple cell lines.
- Artificially reduced ChIP-seq signals are explained by a gain of non-targeted DNA.
- RNA is critical in maintaining the solubility of chromatin during experimentation.

## Introduction

Much of the mammalian genome is transcribed ^1^ with many of these RNAs being retained in the nucleus. RNA that is localised to chromatin can have crucial roles in gene expression and chromatin regulation ^2–7^. RNA has been proposed to serve an architectural role in chromatin organisation with its depletion leading to disruption of both active and repressed chromatin domains ^8–11^. Many chromatin modifiers physically interact with RNA, and direct roles for RNA in their recruitment to target genes have been proposed ^12^. Among the most studied chromatin modifiers in the context of RNA-mediated recruitment is the H3K27me3 methyltransferase Polycomb Repressive Complex 2 (PRC2).

The enzymatic activity and localisation of PRC2 are critical for the repression of cell type-specific genes during embryonic development and are often perturbed in disease ^13–15^. PRC2 interacts with RNA promiscuously ^16^, explained by preferences for prevalent G-tract sequences ^17, 18^. A functional role for PRC2-RNA interactions has been intensely debated, with seemingly opposing models proposed, such as RNA-mediated recruitment of PRC2 or antagonistic interactions (reviewed in ^19, 20^). Further studies have explained aspects of PRC2 recruitment that may occur without RNA involvement ^21–24^. At present, there is ambiguity on how—and to what extent—RNA regulates the chromatin occupancy of PRC2 despite increased usage of perturbation experiments ^18, 25–27^.

For the past number of decades, a common perturbation experiment for the study of RNA-mediated regulation of chromatin has been through RNase treatment ^6, 9, 25, 28–35^. More recently, RNase treatment of chromatin has been used to ascribe functional roles for RNA in the regulation of PRC2 and facultative heterochromatin ^11, 25, 26^. When nuclei are treated with RNase A, ChIP-seq indicates a gain of PRC2 on chromatin ^25^. This data was used to support a model whereby RNA antagonises PRC2 chromatin-binding to dictate the genome-wide localisation patterns of PRC2. Conversely, when chromatin was first isolated from cells and subsequently treated with RNase A, ChIP-seq indicated the depletion of PRC2 subunits from chromatin ^26^. Data that was generated using this method was used to support a model where RNA is required for PRC2 chromatin occupancy through an “RNA bridge” ^26^. These seemingly contradictory observations emphasise a potential problem with the interpretation of ChIP-seq results, if carried out with RNase treatment. The treatment of chromatin with RNAse A during ChIP previously termed either rChIP ^26^ or RNase-ChIP ^35^, is referred to herein as rChIP. While the usage of RNase treatment of chromatin is on the rise, its effect on the properties of chromatin during immunoprecipitation remains unclear.

Here we show that RNase A treatment during chromatin immunoprecipitation (rChIP) reduces PRC2 chromatin occupancy, consistent with previous observations ^26^. However, the reduction in PRC2 ChIP signal is accompanied by a decrease in H3K27me3 ChIP signal genome wide. We track this observation to a global gain of non-targeted chromatin, which takes place during immunoprecipitation if performed with RNase A treatment. As a result, RNase A treatment leads to excess DNA from non-targeted genomic regions. That non-targeted DNA is then sequenced at the expense of DNA from facultative heterochromatin, resulting in the artificial loss of ChIP signals. Collectively, our results point to substantial technical limitations with the usage of RNase treatment of chromatin during ChIP (rChIP) and its application to PRC2. Our results also imply broader limitations of rChIP to other facultative heterochromatin regulators and possibly beyond.

## Results

### The depletion of RNA during chromatin immunoprecipitation is insufficient to change the chromatin occupancy of PRC2

To profile genome-wide PRC2 chromatin occupancy in the context of RNA depletion, we performed ChIP-seq in the presence and absence of RNase A (rChIP) in mouse embryonic stem cells (mESCs). ChIP-seq conditions used in this assay have been successfully applied in many previous publications ^21, 36–41^, and are referred to herein as ‘Condition M’ (see assay conditions in Table 1 and Methods Section). This ChIP protocol was modified to include the addition of RNase A (i.e. rChIP) to the assay after crosslinking and during the immunoprecipitation step, as previously described (^26^ & Figure 1A). Under these rChIP conditions (see Table 1), RNase A treatment did not affect PRC2 chromatin occupancy (Figure 1B-C; hence Conditions M, where ‘M’ stands for “maintenance”). We extended this analysis to RPB1, an RNA PolII subunit associated with active chromatin, and again observed no substantial RNase-dependent changes to its chromatin occupancy (Figure 1B-C). We next confirmed that RNase A effectively eliminates RNA under these ChIP conditions (Condition M; Figure 1D-E). Accordingly, we were also able to detect SUZ12, EZH2 and H3K27me3 enrichment at PRC2 target genes via ChIP-qPCR, either in the presence or absence of RNase A treatment (Figure S1A-C). These results indicate that RNA is dispensable for the retention of PRC2 on chromatin under experimental conditions that enable quality detection of ChIP-seq signals.

**Figure 1.**
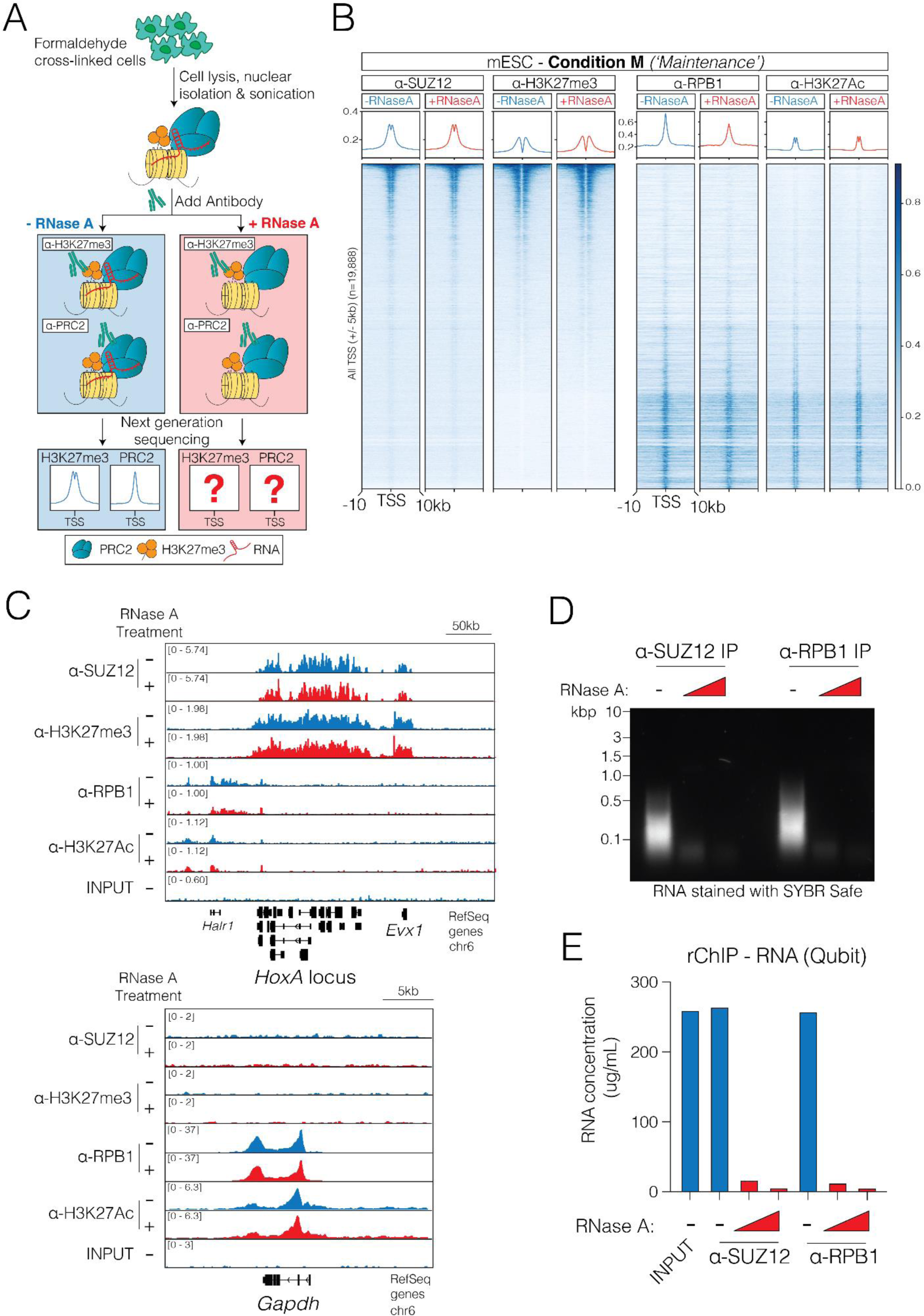
The depletion of RNA during chromatin immunoprecipitation is insufficient to change the chromatin occupancy of PRC2. (A) Schematic illustrating the RNase-ChIP/rChIP assay (see ^26, 43^ & Table 1 for details on differences in ChIP conditions between figures). (B) Heatmap representation and average enrichment profile of SUZ12, H3K27me3, RPB1, H3K27Ac and ChIP-Seq signal at all protein-coding TSS in the presence and absence of RNase A in mouse embryonic stem cells. Data from one representative replicate is shown. ChIP-qPCR from two further independent biological replicates are shown in Figure S1. (C) Genome browser representation of ChIP-seq reads for SUZ12, H3K27me3, RPB1, H3K27Ac and input following RNase A treatment at the *HOXA* gene cluster (repressed) and *GAPDH* gene locus (active). (D) RNA was isolated following RNase A treatment (5 µg/mL or 50 µg/mL) and overnight immunoprecipitation with either SUZ12 or RPB1 antibodies, stained with SYBR Safe and run on an agarose gel, confirming RNase A efficiently removes all RNA in this assay conditions. (E) RNA was isolated following RNase A treatment (5 µg/mL or 50 µg/mL) and overnight immunoprecipitation with either SUZ12 or RPB1 antibodies and RNA concentration was measured on a Qubit RNA High Sensitivity Assay Kit (ThermoFisher Q10211).

**Table 1.** ChIP assay conditions. Specified are variations between the two ChIP conditions that were tested, Condition M and Condition L. Within the Consequence row, in parentheses, indicated are the figures that present data derived using each of the different ChIP conditions. See Methods Section for specific description of the two different ChIP conditions.

| Condition | Condition M | Condition L |
| --- | --- | --- |
| Consequences | Maintenance of PRC2 upon RNA depletion (Figure 1) | Loss of PRC2 upon RNA depletion (Figure 2, 3 & 4 and Long et al. 2020) |
| Crosslinking | 1% formaldehyde (10 min) | 1% formaldehyde (10 min) |
| Isolation of lysate | Nuclei/chromatin | Whole cell |
| Sonication buffer | 100mM NaCl, 33mM Tris-HCl, 1.7% Triton, 0.3% SDS | 0mM NaCl, 50mM Tris HCl, 0.5-1.0% SDS |
| Sonication | 15 min TOTAL "ON" (30s ON/OFF) | 20 min TOTAL "ON" (30s ON/OFF) |
| Cells/ChIP | 2.5x10<sup>6</sup> | 2.5x10<sup>6</sup> |
| IP buffer (final) | 100mM NaCl, 33mM Tris-HCl, 1.67% Triton, 0.33% SDS | 134mM NaCl, 11.7mM Tris-HCl, 0.8% Triton, 0.1-0.2% SDS |

### Loss of PRC2 occupancy following RNA depletion is accompanied by loss of H3K27me3

As we did not observe RNase A-dependent depletion of PRC2 under Condition M (Figure 1), we set out to reproduce the same rChIP conditions that were previously used to identify this phenomenon ^26^ (Table 1, under ‘Condition L’, where ‘L’ stands for “Loss”). Indeed, these ChIP conditions led to an overall reduction in chromatin-bound PRC2 (EZH2) upon RNase A treatment (Figure 2A-B & Figure S2A), as previously reported ^26^. This result is seemingly in agreement with the proposed “RNA-bridging” model for PRC2 chromatin binding ^26^.

**Figure 2.**
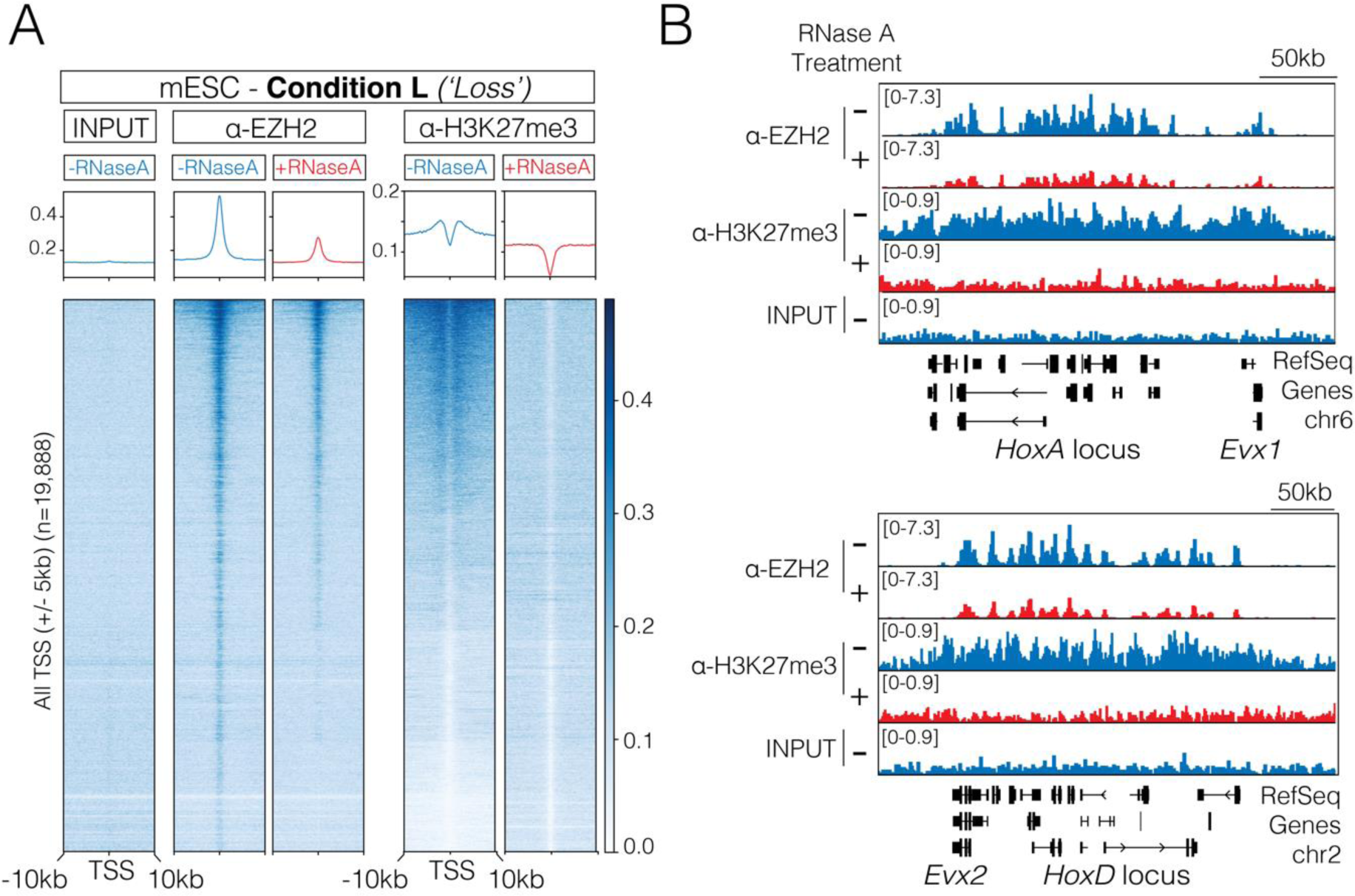
Experimental conditions that favour RNase-dependent depletion of PRC2 chromatin occupancy lead to a concurrent reduction of H3K27me3 ChIP signals. (A) Heatmap representation and average enrichment profile of EZH2 and H3K27me3 ChIP-Seq signal at all protein-coding TSS in the presence and absence of RNase A in mouse embryonic stem cells. One replicate is shown here, and another independent biological replicate is shown in Figure S2. (B) Genome browser representation of ChIP-Seq reads for EZH2 and H3K27me3 following RNase A treatment at both the *HOXA* and *HOXD* gene clusters.

To further challenge the rChIP assay under Condition L, we next probed for H3K27me3 as a control (see Figure 2A-B). H3K27me3 was selected as a negative control because we had no reason to assume that an RNA bridge would be present between H3K27me3 and chromatin. Yet, the RNA-dependent reduction in PRC2 was accompanied by a concurrent loss in H3K27me3 rChIP-seq signal (Figure 2A-B & Figure S2A). This apparent loss of facultative heterochromatin following RNA depletion is consistent with another previous study that observed decreases in H3K27me3, H3K9me3 as well as CTCF upon RNase A treatment in a CUT&RUN assay ^11^. Hence, results thus far suggest that RNase treatment under Condition L leads to the apparent loss of ChIP-seq signals from facultative heterochromatin, whether probing for PRC2 (ɑ-EZH2) or repressive chromatin itself (ɑ-H3K27me3).

### RNA depletion leads to the apparent loss of H3K27me3 rChIP signals in human induced pluripotent stem cells (iPS) and cancer cells

All data presented to this point was generated in mESCs. Yet, previously reported rChIP experiments using antibodies for PRC2 subunits were carried out in human induced pluripotent stem cells (iPSCs) ^26^. Therefore, before testing the hypothesis of an rChIP-associated artefact, we first wished to determine if the previously published rChIP conditions (Condition L from ^26^) lead to the apparent depletion of H3K27me3 in human iPSC cells. We, therefore, performed the rChIP assay in iPSCs as previously published for human iPSC (Condition L ^26^; Figure 3).

**Figure 3.**
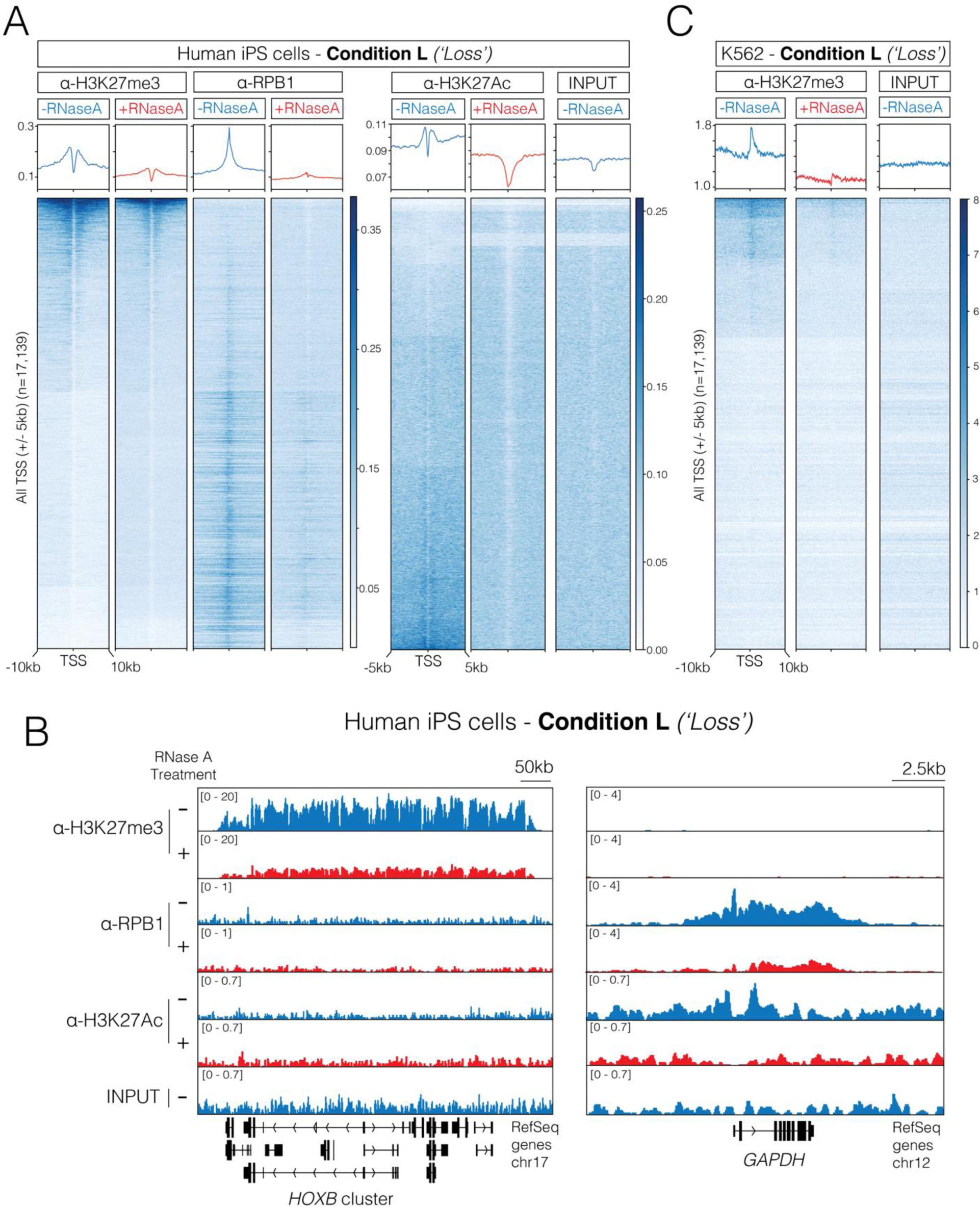
RNA depletion leads to the apparent loss of ChIP-signals from facultative heterochromatin in human iPS and cancer cells. (A) Heatmap representation and average enrichment profiles of H3K27me3, RPB1, H3K27Ac and input ChIP-seq signal at all human protein-coding TSS in the presence and absence of RNase A in human induced pluripotent stem cells (iPS cells). Data illustrates a loss of both facultative heterochromatin (H3K27me3) and euchromatin (RPB1 & H3K27Ac) following RNA depletion. One replicate is shown here, and another independent biological replicate is shown in Figure S3. (B) Genome browser representation of ChIP-seq reads for H3K27me3, RPB1, H3K27Ac and input following RNase A treatment at the *HOXB* gene cluster (repressed) and *GAPDH* gene locus (active) in human iPS cells. (C) Heatmap representation and average enrichment profile of H3K27me3 and input ChIP-seq signal at all human protein-coding TSS in the presence and absence of RNase A in K562 human cancer cells. Experiments in this figure were carried out under Condition L (see Table 1 for details on differences in ChIP conditions between figures). Data from one replicate is shown.

Once more we observed a reduction in the H3K27me3 signal upon RNase A treatment (Figure 3A-B & Figure S3A-B), in agreement with our results using mESC (Figure 2). We also observed this phenomenon in lineage-committed human cancer cells (K562) (Figure 3C & Figure S3C). This suggested that RNA depletion is leading to a global loss of rChIP signals from facultative heterochromatin. As expected, the loss of H3K27me3 with RNase A treatment in iPSCs was accompanied by a loss of PRC2 (SUZ12 & EZH2; Figure S3D-E). Furthermore, we also saw a decrease in RPB1 and H3K27Ac rChIP-seq signals in iPSCs (Figure 3A-B & Figure S3A-B), suggesting that an artificial reduction of ChIP-seq signals also occurred in euchromatin. Observations thus far indicate that ChIP signals of both chromatin-bound factors and histone marks can be disrupted by RNA depletion.

### RNA depletion during ChIP leads to a global gain of non-targeted DNA

Data thus far indicate that RNase treatment during ChIP under Condition L (Table 1) leads to the loss of ChIP signals from facultative heterochromatin and possibly euchromatin (Figure 2 & Figure 3). Yet, there is no obvious biological explanation for the loss of H3K27me3 after RNA depletion, as RNase treatment was carried out after chromatin was crosslinked and isolated from cells, and hence cannot be due to the loss of PRC2 methyltransferase activity. Therefore, we were next determined to test the hypothesis of an experimental artefact. To this end, we performed ChIP-qPCR for H3K27me3, H3K27Ac and SUZ12 in the presence and absence of RNase A in human iPSCs under Condition L. ChIP-qPCR was selected for this analysis given its ability to directly quantify the IP sample with respect to the input, without reliance on genome-wide normalisation effects.

As expected, in the absence of RNase A treatment, ChIP-qPCR identified H3K27me3 and SUZ12 at a repressed gene (*HOXA10*) and H3K27Ac on active genes (*GAPDH* and *CCNA2*; Figure 4A & Figure S4A, blue bars). But strikingly, RNase A treatment led to an increment of the ChIP-qPCR signal at all loci for all the antibodies tested (Figure 4A & Figure S4A, red bars). This was inconsistent with the ChIP-seq results we obtained with RNase A treatment, which reported the loss of ChIP-seq signal under the same conditions (Condition L; Figures 2 & 3).

**Figure 4.**
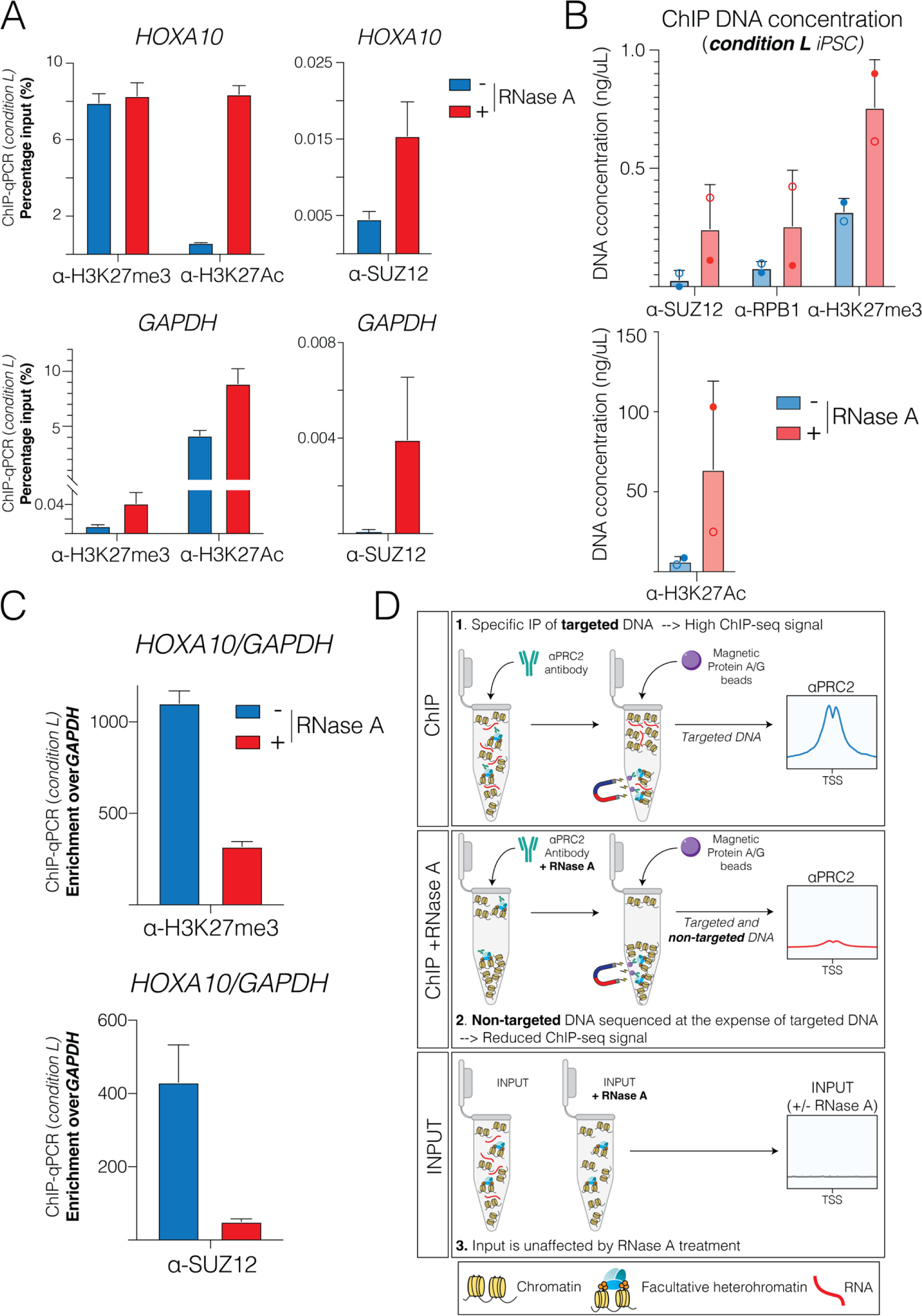
RNA depletion during ChIP leads to a global gain of non-targeted chromatin, which artificially reduces ChIP signals upon data normalisation. (A) ChIP-qPCR analyses of H3K27me3, H3K27Ac and SUZ12 at a representative repressed PRC2 target TSS (*HOXA10*; repressed gene) and active TSS (*GAPDH*; active gene) following RNase A treatment in human iPS cells. Enrichment is presented as a percentage of input DNA. One replicate is shown here, and an independent biological replicate is shown in Figure S4C-D. (B) The concentration of DNA immunoprecipitated with the antibodies shown was measured on a Qubit dsDNA High Sensitivity Assay Kit (ThermoFisher Q32854). Qubit readings from two independent rChIP replicates are shown (solid circles: replicate 1; empty circles: replicate 2). (C) ChIP-qPCR analyses from (A) presented as enrichment at the repressed *HOXA10* TSS normalised to a negative control TSS *GAPDH*. (D) Schematic illustration of the potential mechanism behind the loss of facultative heterochromatin after RNase A treatment in a ChIP assay: The depletion of RNA during immunoprecipitation reduces the solubility properties of chromatin, leading to the accumulation of both targeted and non-targeted chromatin aggregates in the IP sample. After ChIP and data normalisation, the DNA from untargeted chromatin artificially reduces the signals from facultative heterochromatin. Experiments in this figure were carried out under Condition L (see Table 1 for details on differences in ChIP conditions between figures).

Hence, results so far are seemingly contradictory: RNase A treatment under Condition L leads to a gain of ChIP-qPCR signal (Figure 4) but a loss of ChIP-seq signal (Figures 2 & 3). The simplest explanation that consolidates all these observations is a global gain of non-targeted DNA. This is emphasised by the observation that RNase A treatment led to the unexpected gain of H3K27me3 ChIP-qPCR signal at active genes as well as the unexpected gain of H3K27Ac signals at repressed genes (Figure 4A & Figure S4A). To directly test for an RNase A-dependent gain of DNA, we quantified the amount of total DNA that was obtained after ChIP, in the presence and absence of RNase A. Indeed, RNase A treatment increased the total amount of immunoprecipitated DNA by 2- to 10-fold with respect to the untreated chromatin, with some variations between replicates and antibodies used (Figure 4B & Figure S4B). We also confirmed this increase in immunoprecipitated DNA does not occur when RNA is depleted from chromatin isolated from iPSCs under Condition M (Figure S4C). These results strongly indicate that RNase treatment under Condition L does not lead to the depletion of PRC2-bound chromatin, but it in fact leads to a gain of non-targeted chromatin.

### Loss of ChIP signals after RNase treatment as a normalisation artefact

Data thus far indicates that RNase A treatment leads to the depletion of ChIP signals, for PRC2 subunits and H3K27me3 (Figure 2 & 3), but also to a global gain of non-targeted DNA (Figure 4A-B & Figure S4A-B). The simplest explanation for these observations is that the apparent depletion of rChIP-seq signals herein (Figure 2 & 3) and elsewhere ^26^ resulted from a normalisation artefact. Such an artefact could occur if DNA at target genes is normalised against DNA elsewhere. This is a common practice with ChIP-seq normalisation, where signals are normalised against the total aligned reads. The artificial reduction of ChIP-seq signals after RNase A treatment is expected to persist under most types of commonly used normalisation approaches. This is because the loss of ChIP-seq signal is, in fact, attributed to a gain in the background: DNA from non-targeted chromatin is sequenced at the expense of DNA from targeted chromatin. As a demonstration of this effect, when rChIP-qPCR signals from the repressed *HOXA10* gene are normalised against the active *GAPDH or CCNA2* TSS, both the SUZ12 and H3K27me3 enrichments appeared reduced after RNase A treatment (Figure 4C and Figure S4D). Conversely, we did not observe an artificial reduction of H3K27me3 if the iPSC chromatin was treated under Condition M (Figure S4E-F).

Collectively, our results indicate that the complete depletion of RNA from chromatin is insufficient to alter PRC2 chromatin occupancy (Figure 1; Condition M in Table 1). Yet, rChIP conditions that lead to the apparent depletion of PRC2 from chromatin also lead to the depletion of H3K27me3 (Figure 2 & 3, and Condition L in Table 1). All these observations can be explained by a global gain of non-targeted DNA after RNase A treatment (Figure 4A,C & Figure S4A,C), which artificially leads to the reduction of ChIP-seq signals as a result of a normalisation artefact (Figure 4B,D & S4B).

## Discussion

RNA has been shown to play an architectural role in chromatin organisation and to be essential for the integrity of both constitutive and facultative heterochromatin^2, 6, 8, 9, 11, 25, 28–30^. While RNase treatment of chromatin has been central in these studies, we are only beginning to understand how this treatment affects the biophysical properties of chromatin. In practice, RNA degradation may reduce the solubility of chromatin under some ChIP conditions and buffers. Indeed, the observation that RNase treatment alters the structural properties of chromatin and leads to the formation of “massive clumps” has been reported at least since the 1980s ^6, 28^. Our analyses point to challenges in interpreting ChIP results if performed with RNase A treatment.

Our observations imply that a previous proposal of an “RNA bridge” between PRC2 and chromatin ^26^ has been supported based on an experimental artefact: RNase treatment under certain ChIP conditions (Condition L; Figure 2, Figure 3 & ^26^) leads to a gain of non-targeted DNA (Figure 4 & Figure S4). We speculate that this could be the result of reduced chromatin solubility in the absence of RNA, which in turn leads to more chromatin precipitating with the beads (Figure 4D). In agreement with this, two independent parallel studies confirmed this phenomenon: Hall Hickman and Jenner demonstrated that precipitated chromatin could be visually detected on the beads after RNase A treatment ^42^ and Long, Hwang and colleagues quantified excess DNA in the IP sample after RNase A treatment ^43^.

RNase A treatment has been shown to increase the tendency of chromatin to precipitate, especially when carried in the presence of a low concentration of monovalent salt ^44^. This salt effect could be a potential explanation for why Condition L leads to RNase A-dependent depletion of ChIP signals (Figure 2-4 and ^26^) while Condition M does not (Figure 1): Condition L (and ^26^) contains less monovalent salt in the sonication buffer than Condition M (Table 1). The salt effect could have been further amplified in the case of Condition L, where whole cell lysates were isolated as opposed to nuclear lysates in Condition M (Table 1). Specifically, the lack of a nuclear isolation step in Condition L could have increased the total amount of RNA during the sonication and IP, with respect to Condition M which included nuclear isolation (Table 1). The excess RNA under Condition L could have solubilised chromatin, despite the low salt in the sonication buffer, possibly given its negative charge. Consistent with a potential role for RNA negative charges in solubilising chromatin, a parallel study demonstrated that the anionic polymer Poly-L-glutamic acid (PGA) is capable of restoring ChIP signals after RNase treatment ^42^. In further support of this model, another parallel independent study demonstrated that the catalytic activity of RNase A is required for the depletion of ChIP-seq signals ^43^. Hence, as soon as RNA is eliminated under Condition L, the low salt may have caused chromatin to precipitate. Once non-targeted chromatin is precipitated with the beads, ChIP-seq signals are reduced because excess DNA from non-targeted chromatin is sequenced at the expense of DNA from targeted chromatin.

The original study ^26^ did not include H3K27me3 as a control for rChIP, while we did (Figure 1-4). Yet, they did use antibodies for the euchromatic factors RBP1 and TBP, which were not sensitive to RNase A digestion ^26^. We were unable to reproduce these results: in our hands, RNase A treatment also led to the reduction of ChIP-seq signals and gain of DNA in the case of RBP1 (Figures 3 and 4). RNase A treatment increased the amount of DNA that precipitated with all antibodies tested, with some variations between different antibodies and independent replicates (Figure 4B). Variations in the amount of immunoprecipitated DNA were accompanied by variations in the rChIP-seq signals (compare Figure 2A-B and Figure S2A). These observations suggest suboptimal robustness of rChIP under Conditions L, in agreement with recent data from an independent study ^43^. The global gain of DNA after ChIP with RNase treatment (Figure 4 & Figure S4) does not fit well with the disruption of RNA bridges between the targeted factor to chromatin.

Our results do not reflect poorly on previous studies that used RNase treatment for applications other than ChIP. It is also important to note that our work cannot absolutely exclude the possibility of an RNA bridge between PRC2 to chromatin. Yet, our results firmly support two conclusions: (i) RNA is dispensable for the PRC2 chromatin occupancy after cells were already crosslinked and chromatin has been isolated (Figure 1); and (ii) the “RNA bridging” model for PRC2 chromatin occupancy ^26^ has been supported based on an experimental artefact (Figure 4). RNase treatment will likely remain useful for the study of RNA-mediated regulation of chromatin modifiers as long as the global biophysical effects of RNA degradation are considered and controlled for.

## Limitations of the study

Our study was designed to determine if RNase A treatment under Condition L reports for real changes in PRC2 chromatin occupancy or rather if it leads to the artificial depletion of ChIP-seq signals. Our study was not designed to identify the responsible determinant within Condition L, yet, biochemical analysis by Dueva et al.^44^ points to the concentration of monovalent salt as a key determinant of chromatin solubility after RNase A treatment. Accordingly, Condition L is characterised by a low concentration of monovalent salt compared to Condition M.

We did not characterise Condition M as thoroughly as we did with Condition L. This is because our study was not designed to exclude an RNA bridge between PRC2 and chromatin, but rather to examine evidence previously used to support it. Future studies would have to demonstrate the robustness of rChIP under Condition M or other conditions, if they are to be used for discovery research. Demonstration of rChIP robustness would be important, given the variations in the amount of the immunoprecipitated DNA on different days under Condition L and while using different antibodies. Experiments herein may serve to guide this process.

## Acknowledgements

We are grateful to the members of the Davidovich lab for helpful discussions in relation to this work. We thank Richard Jenner, Tom Cech and John Rinn for discussions of our work. We would also like to thank the support from the MASSIVE HPC facility. Q.Z. is supported by Investigator Grant EL1 (APP1196365) from National Health and Medical Research Council (NHMRC). C.D. is an EMBL-Australia Group Leader and a Sylvia and Charles Viertel Senior Medical Research Fellow and acknowledges support from the NHMRC (GNT1184637, GNT2011767 & GNT2020900).

## Author Contributions

E.H., Q.Z. and C.D. conceived the project and designed the experiments. E.H. and Q.Z. carried out all lab-based experiments. E.H., Q.Z. and E.H.G. performed the bioinformatic analyses of ChIP-seq datasets. G.S, M.B, V.P, P.P.D & J.M.P provided expertise on the culturing of human and mouse pluripotent stem cells. S.C.A cultured mESCs for rChIP experiments in Figure 2. E.H. and C.D. co-wrote the manuscript. C.D. supervised.

## Data Availability

All next-generation sequencing datasets have been deposited and can be accessed via accession number GSE240079.

## Materials and Methods

### Cell Culture

Mouse embryonic stem cells (mESCs) were grown on gelatinised culture dishes in DMEM supplemented with 10 % FBS (Scientifix; FBSAU-2007A), 1 % (v/v) penicillin-streptomycin (Thermo Scientific #15140122), 50 µM beta-mercaptoethanol (Millipore; ES-007-E), 1:100 non-essential amino acids, 1:100 sodium pyruvate, 1:100 GlutaMax (all GIBCO) and ESGRO leukemia inhibitory factor (LIF) (Millipore; ESG1107; 10,000x).

K562 cells were cultured in RPMI-1640 (Merck #R8758) growth medium supplemented with 10 % FBS (Cellsera AU-FBS/SF) and 1 % (v/v) penicillin-streptomycin (Thermo Scientific #15140122) and were incubated at 37 °C with 5 % CO_2_. K562 cells were acquired from ATCC.

Human induced pluripotent stem cells (iPS) were derived from HDFn (neonatal human dermal fibroblasts) according to ^45^. HDFn iPS cells were maintained in a feeder-free system on vitronectin (VTN-N; Gibco; A31804) coated tissue culture plastics in Essential 8 Flex medium (Gibco; A2858501). Cells were incubated at 37 °C and 5 % CO_2_ and passaged every 2-3 days using 0.5mM EDTA. Human iPS cells were obtained from the Jose Polo Lab. All cell lines were tested periodically for mycoplasma contamination.

### Formaldehyde crosslinking for Chromatin immunoprecipitation (ChIP)

Mouse ESC, K562 or human iPS cells were collected, counted, and washed once with PBS before crosslinking for 10 minutes with PBS containing 1 % formaldehyde (Sigma). Cells were crosslinked in 15 mL falcon tubes at a density of ∼5×10^6^ cells/mL. Crosslinking was quenched with 0.125 M Glycine prior to two PBS washes. Crosslinked cells were then snap-frozen on liquid nitrogen and stored at −80°C before proceeding with the rest of the ChIP assay.

### ChIP conditions resulting in no changes to the PRC2 occupancy following RNase A treatment (Condition M)

ChIP assay was performed as described previously ^21, 36, 39, 40^. Crosslinked cells were thawed on ice and lysed in 5 mL of SDS-Lysis buffer (100 mM NaCl, 50 mM Tris pH 8.1, 5 mM EDTA pH 8.0, 0.5 % SDS, 1X protease inhibitor cocktail [Sigma; P8340]). Nuclei and chromatin were pelleted by centrifugation at 1200 rpm for 6 min at room temperature. The supernatant was then discarded, and the pellet was resuspended in 2 mL of ChIP buffer (33 mM Tris-HCl pH 8, 100 mM NaCl, 5 mM EDTA, 0.33% SDS, 1.67% Triton X-100, 1X protease inhibitor cocktail [Sigma; P8340]). Chromatin was sheared to approximately 200 bp to 500 bp fragments by sonication using a Bioruptor Plus (Diagenode) at high power. Total sonication time (i.e., cumulative “on” time) was 15min with 30 sec on/off pulses. Sonicated chromatin was incubated overnight with antibody while rotating at 4 °C. Prior to overnight incubation with antibodies, an input sample was taken (1 %). RNase A (DNase and protease-free Thermo Fisher Scientific; #EN0531) was added along with the antibody to the plus RNase A samples at a final concentration of 5-50 µg/mL. Extract from 2.5×10^6^ cells were used per ChIP (+/- RNase A). The following morning, samples were clarified by centrifugation at 20,000 g for 20 min at 4 °C. Following clarification, the chromatin was incubated for 3 hours with Protein G Dynabeads (ThermoFisher; 10004D; 50 µL beads were used per ChIP). After incubation, the beads were washed three times in Mixed Micelle Buffer (150 mM NaCl, 20 mM Tris pH 8.1, 5 mM EDTA pH 8.0, 5.2 % Sucrose, 1 % Triton X-100, 0.2 % SDS), twice with Buffer 500 (0.1 % Sodium Deoxycholate, 1 mM EDTA pH 8.0, 50 mM HEPES pH 7.5, 1 % TritonX-100), twice with LiCl detergent wash (0.5 % Sodium Deoxycholate, 1 mM EDTA pH 8.0, 250 mM LiCl, 0.5 % NP-40,10 mM Tris pH 8.0) and finally, one wash with TE. All the washes were performed at 4°C. Immunoprecipitated material was eluted from the beads with 100 µL elution buffer (0.1 M NaHCO3, 1 % SDS) while shaking for 1 hour at 65 °C. The supernatant was retained and incubated overnight at 65 °C while shaking to reverse the crosslinks. The eluted material was then subjected to RNase A (DNase and protease-free Thermo Fisher Scientific; #EN0531) and Proteinase K (ThermoFisher; #EO0491) treatments prior to DNA clean up (QIAGEN MinElute PCR Purification Kit Cat#28004). ChIP enrichments were analysed by qPCR using the QuantiNova SYBR Green PCR kit (Qiagen; 208054).

When testing RNase A efficiency during the rChIP assay, 10% of lysate was removed following overnight immunoprecipitation and RNase A treatment. Protein was removed with Proteinase K (ThermoFisher; #EO0491) and crosslinks reversed for 2hrs at 55 °C. RNA was extracted with 1 mL TRIzol and 200 µL chloroform, inverted several times and centrifuged at 20,000g for 20 min at 4 °C. The aqueous layer was then passed through a QIAGEN RNeasy minElute column. Following one wash with RWT buffer DNA was digested on the column with DNase I for 15 min at room temperature. Following washes in RPE buffer RNA was eluted in RNase free water and analysed by electrophoresis on an agarose gel and quantified using the Qubit RNA High Sensitivity Assay Kit (ThermoFisher Q32852).

### ChIP conditions resulting in RNase-dependent loss of PRC2 occupancy (rChIP; Condition L)

ChIP assay with RNase A treatment was performed based on a previous study ^26^. Specifically, crosslinked cells were thawed on ice, lysed in 2 mL lysis buffer (50 mM Tris-Cl pH 8.1, 10 mM EDTA, 0.5-1.0 % SDS, plus 1X protease inhibitor cocktail [Sigma; P8340]) and incubated on ice for 10 mins. Chromatin was sheared to approximately 200bp-500bp fragments by sonication using a Bioruptor Plus (Diagenode) at high power. Total sonication time (e.g., total “on” time) was 20 mins with 30 sec on/off pulses. After sonication, the lysate was cleared by centrifugation at 16,300 g for 10 min at 4 °C. The supernatant was stored at −80°C or used directly in the ChIP assay. Lysate was diluted 1:5 in IP buffer (16.7 mM Tris-HCl pH 8.1, 1.2 mM EDTA, 167 mM NaCl, 1% Triton X-100 and 1X protease inhibitor cocktail [Sigma; P8340]) and pre-cleared with 50 µL protein G Dynabeads (ThermoFisher; 10004D) for 1 hour at 4 °C. The sonicated lysate was incubated overnight with antibody while rotating at 4 °C. Prior to overnight incubation with the antibodies, an input sample was taken (1%). RNase A (DNase and protease-free Thermo Fisher Scientific; #EN0531) was added along with the antibody to the plus RNase A samples at a final concentration of 50 µg/mL. Extracts from 2.5×10^6^ cells were used per ChIP (+/- RNase A). The Next morning, Protein G Dynabeads were equilibrated in IP buffer: 25 µL beads were added to each ChIP and incubated for 1 hour at 4°C. After incubation, beads were washed twice in low salt buffer (20 mM Tris-HCl pH 8.0, 2 mM EDTA, 150 mM NaCl, 0.1 % SDS, 1 % Triton x-100), twice with high salt buffer (20 mM Tris-HCl pH 8.0, 2 mM EDTA, 500 mM NaCl, 0.1 % SDS, 1% Triton x-100), twice in LiCl buffer (10 mM Tris-HCl pH 8.0, 1 mM EDTA, 250 mM LiCl, 1% Sodium Deoxycholate (w/v), 1% NP-40 substitute) followed by one final wash in TE buffer (10 mM Tris pH 8.0, 1 mM EDTA). All the washes were performed at 4 °C. Immunoprecipitated material was eluted from the beads with 120 µL elution buffer (0.1 M NaHCO3, 1% SDS) for 20 min at room temperature. The supernatant was retained and 200 mM NaCl was added and incubated along with input samples overnight at 65 °C while shaking to reverse the crosslinks. The eluted material was then subjected to RNase A (DNase and protease-free Thermo Fisher Scientific; #EN0531) and Proteinase K (ThermoFisher; #EO0491) treatments prior to DNA clean up (QIAGEN MinElute PCR Purification Kit Cat#28004). ChIP enrichments were analysed by qPCR using the QuantiNova SYBR Green PCR kit (Qiagen; 208054).

### Antibodies

Anti-EZH2 (Cell Signalling; #5246), 8 µg per ChIP, Anti-SUZ12 (Cell Signalling; #3737), 0.3 µg per ChIP. Anti-H3K27me3 (Cell Signalling; C36B11), 5 µg per ChIP. Anti-H3K27Ac (Abcam; ab4729), 5 µg per ChIP. Anti-RPB1 (NTD) (Cell Signalling; #14958) 1 µg per ChIP.

### ChIP-qPCR primer sequences (5’-3’)

m*Pax3*-F - AGAAAGGCGGAAAGAGATTAGGAC

m*Pax3*-R - GAGCCAGTGAGGATGGAAAAAG

m*Sox1*-F - TTTGCACAGTTCAGCCCTGAGTGA

m*Sox1*-R - GGTGCACAAACCACTTGCCAAAGA

m*T*-F - TCCGCAGAGTGACCCTTTTTC

m*T*-R - TACCCAACAGCCACCTTCACTTC

m*Gapdh*-F - AGTGTGCACCAAGGACATCCAG

m*Gapdh*-R - CCCATTTTACTCGGGAAGCAG

h*HOXA10*-F - CCCGAGCTGATGAGCGAGTC

h*HOXA10*-R - GCCAAATTATCCCACAACAATGTC

h*GAPDH*-F - TACTAG CGGTTTTACGGGCG

h*GAPDH*-R - TCGAACAGGAGGAGCAGAGAGCGA

h*CCNA2*-F - TGACGTCATTCAAGGCGA

h*CCNA2*-R - GCTCAGTTTCCTTTGGTTTAC

### ChIP-Seq library preparation

Following ChIP (+/- RNase A) experiments, the precipitated DNA was quantified using the Qubit dsDNA High Sensitivity Assay Kit (ThermoFisher Q32854). A total of 1-10 ng of DNA from each ChIP experiment was used for library preparation using the NEBNext Ultra II DNA Library Kit for Illumina (E7645) and NEBNext Multiplex Oligos for Illumina (Set#1 & #2; E7335 & E7500). Following adaptor ligation, DNA was PCR amplified for 4-10 cycles, depending on the amount of the input DNA. DNA purification was then performed using NEBNext Sample Purification Beads (E7767S). The quality and size distributions of DNA libraries were ascertained on a TapeStation (Agilent) using High Sensitivity D1000 ScreenTape assay reagents (Agilent; 5067-5585). The resulting libraries were then used for cluster generation and sequencing on an Illumina-based platform with a 150 bp paired-end read format (2-3×10^7^ reads per sample) (GeneWiz/Azenta).

### Bioinformatic analysis of ChIP-Seq (+/- RNase A) datasets

ChIP-Seq reads were aligned to a reference genome (mm10 or hg38) using Bowtie2 with default parameters ^46^. SAMtools was used to convert SAM files to BAM files and to remove duplicate aligned reads ^47^. Bigwig files were generated using bamCoverge from the deepTools suite (version 3.1.3) ^48^ with a bin size of 10 and scaled using CPM (counts per million) normalisation. All transcription start sites (TSS) for the mm10 (n=19,888) and hg38 (n=17,139) builds of the mouse and human genomes were defined and annotated as unique protein-coding TSS +/-5kb (downloaded from ENSEMBL biomart). Heatmaps were generated with these TSS annotation files using the computeMatrix and plotHeatmap tools from the deepTools suite ^48^. Peak calling was performed using MACS2 ^49^ with a p-value cut-off of 1×10^-4^.

---

**Figure S1.**
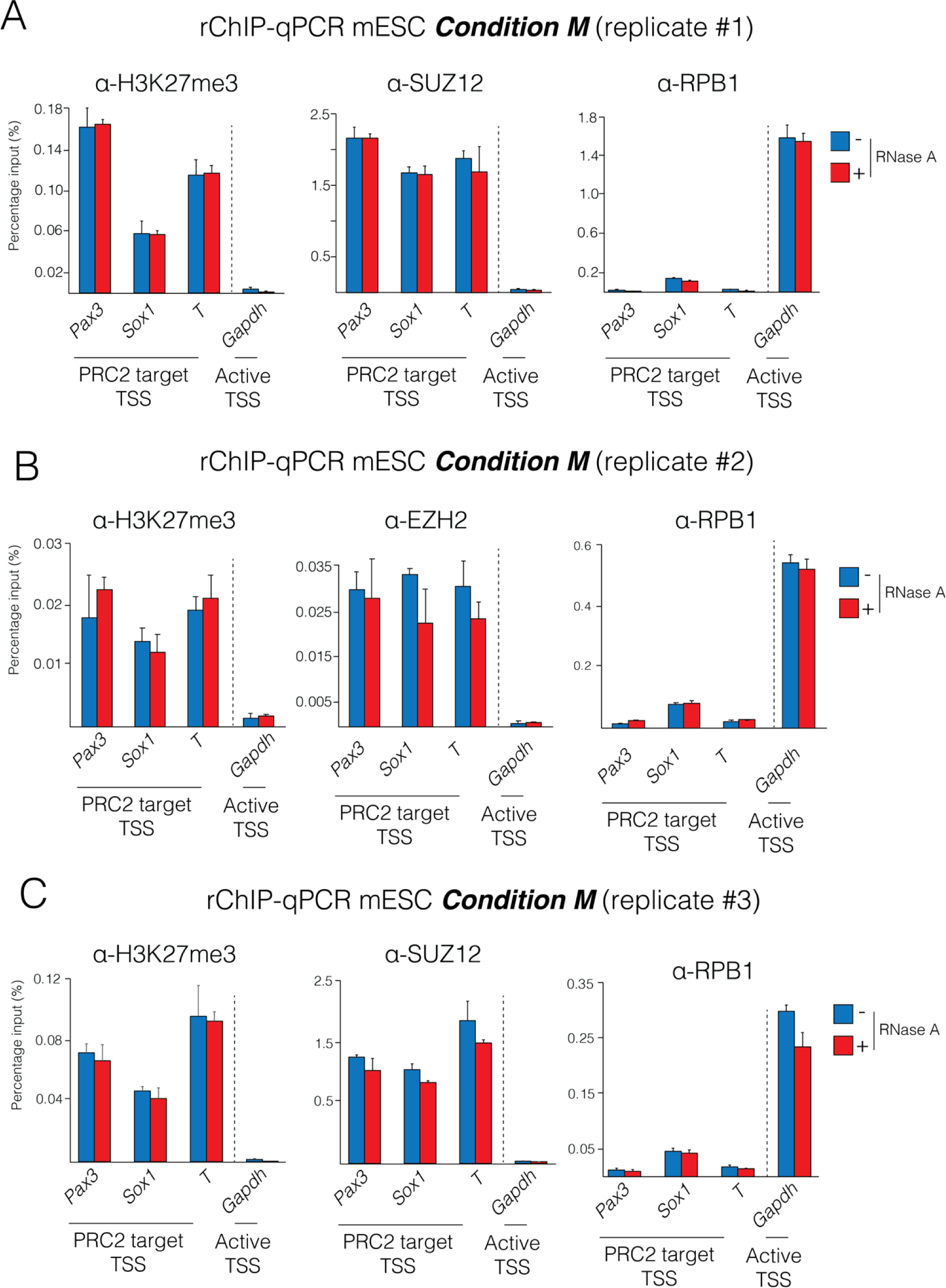
The depletion of RNA during chromatin immunoprecipitation is insufficient to change the chromatin occupancy of PRC2. (A), (B) & (C) rChIP-qPCR analyses from three independent biological replicates for H3K27me3, SUZ12 or EZH2 and RPB1 at representative repressed PRC2 target TSS (*Pax3*, *Sox1* & *T*) and active gene TSS (*Gapdh*) following RNase A treatment (50 µg/mL treatment shown). Experiments in this figure were carried out under Condition M (see Table 1 for details on differences in ChIP conditions between figures). Error bars indicate standard deviation of qPCR triplicates. Replicate #1 is from the same rChIP-seq experiment shown in Figure 1, while replicates #2 and #3 are independent biological replicates.

**Figure S2.**
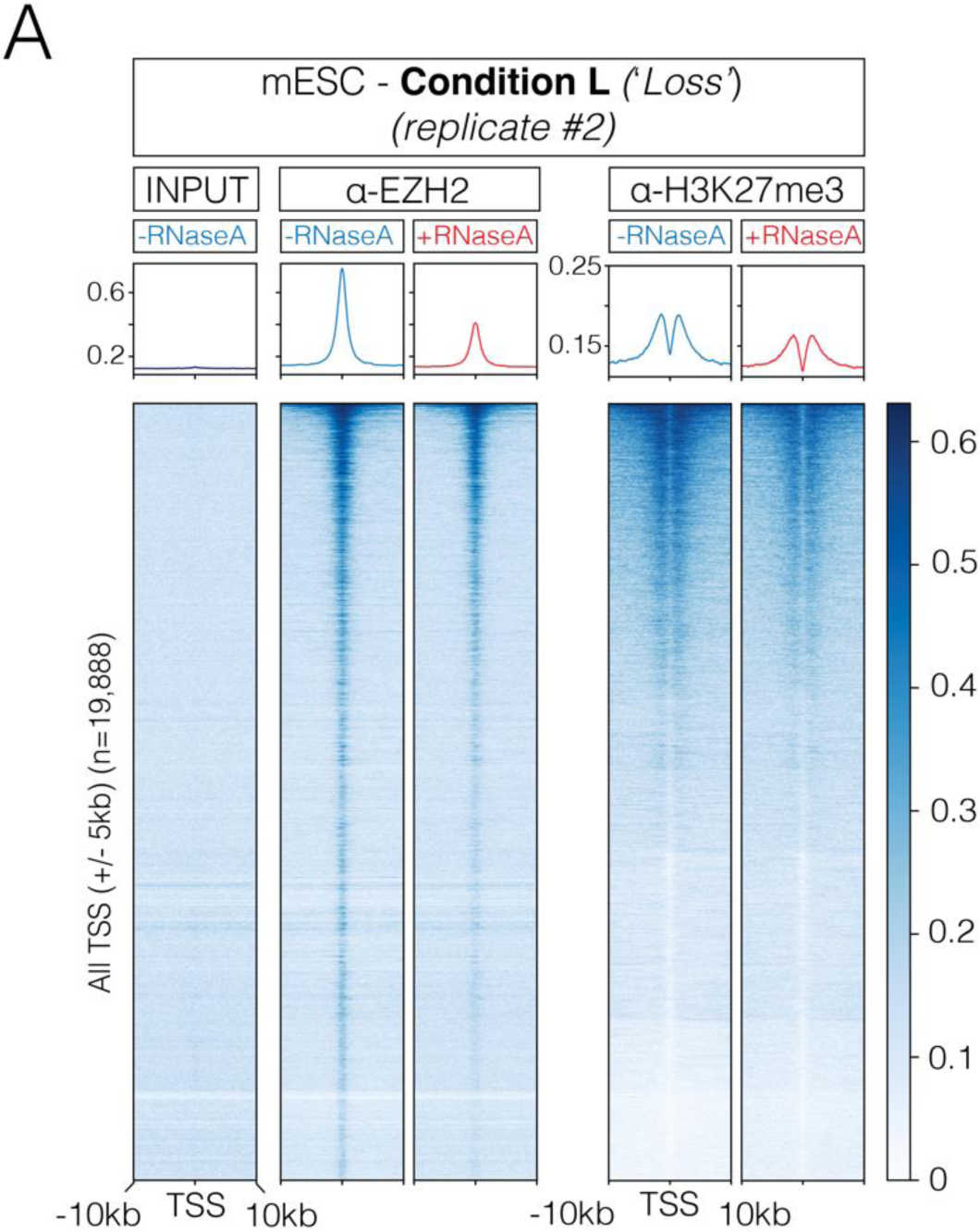
Experimental conditions that favour RNase-dependent depletion of PRC2 chromatin occupancy lead to a concurrent reduction of H3K27me3 ChIP signals. (A) Heatmap representation and average enrichment profile of EZH2 and H3K27me3 ChIP-Seq signal at all protein-coding TSS (n=19,888) in the presence and absence of RNase A in mouse embryonic stem cells. The data shown here is an independent replicate to the data in Figure 1A, which was carried on a different day. Experiments in this figure were carried out under Condition L (see Table 1 for details on differences in ChIP conditions between figures).

**Figure S3.**
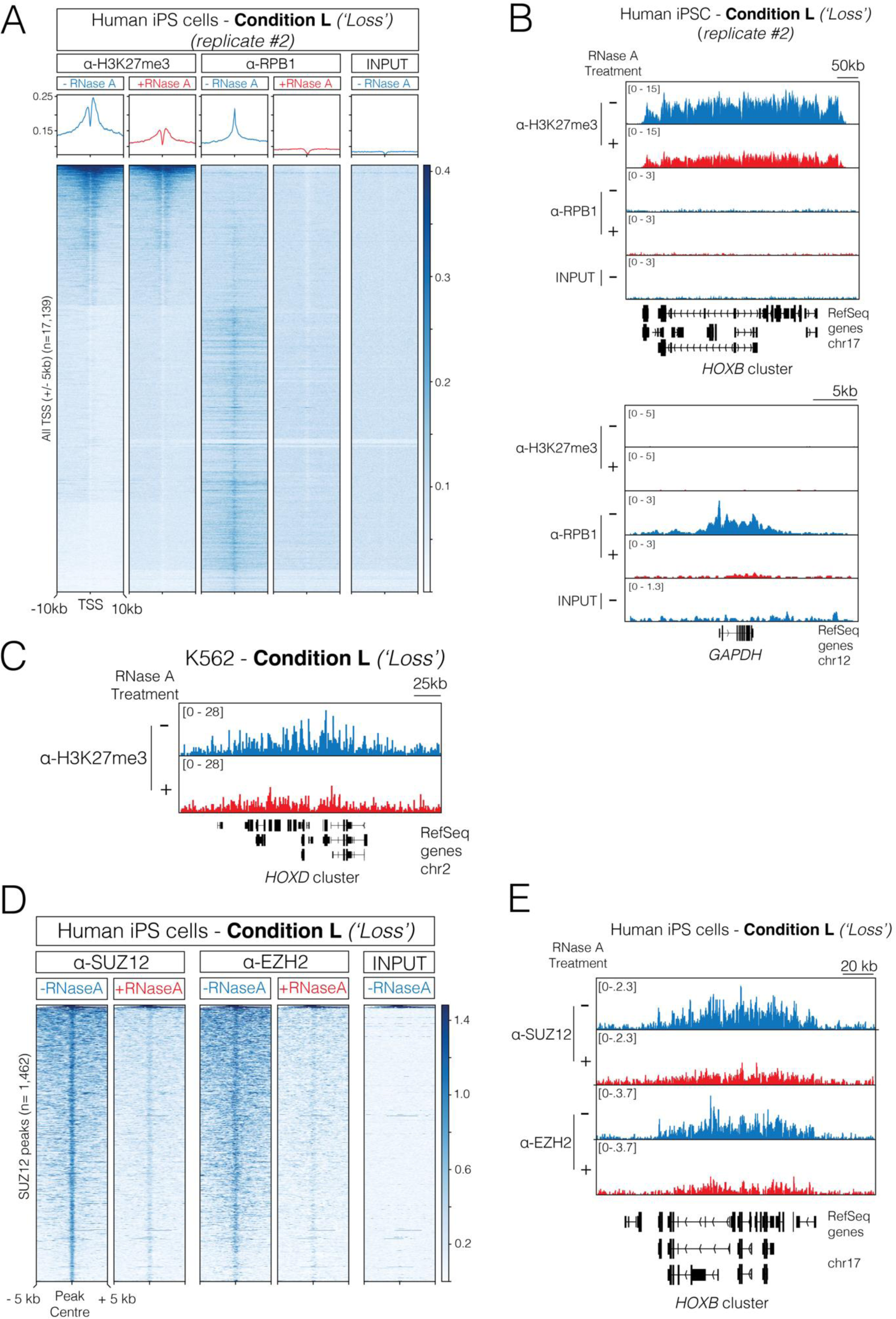
RNA depletion leads to the apparent loss of ChIP-signal from facultative heterochromatin in human iPS and cancer cells. (A) Heatmap representation and average enrichment profile of H3K27me3, RPB1 and input ChIP-Seq signal at all human protein-coding TSS (n=17,139) in the presence and absence of RNase A in human induced pluripotent stem cells (iPS cells). Data shown here is from an independent replicate of the experiment presented in Figure 3A-B. (B) Genome browser representation of ChIP-Seq reads for H3K27me3, RPB1 and input following RNase A treatment at the *HOXB* gene cluster (repressed) and *GAPDH* gene locus (active). Data shown here is from an independent replicate of the experiment presented in Figure 3A-B. (C) Genome browser representation of ChIP-Seq reads for H3K27me3 following RNase A treatment at the *HOXD* cluster in K562 cells. Experiments in this figure were carried out under Condition L (see Table 1 for details on differences in ChIP conditions between figures). (D) Heatmap representation of SUZ12 and EZH2 ChIP-Seq signal at all SUZ12 peaks (n=1,462) in the presence and absence of RNase A in human induced pluripotent stem cells (iPS cells). (E) Genome browser representation of ChIP-Seq reads in the presence and absence of RNase A for SUZ12 and EZH2 ChIP-Seq signal at the *HOXB* gene cluster (repressed).

**Figure S4.**
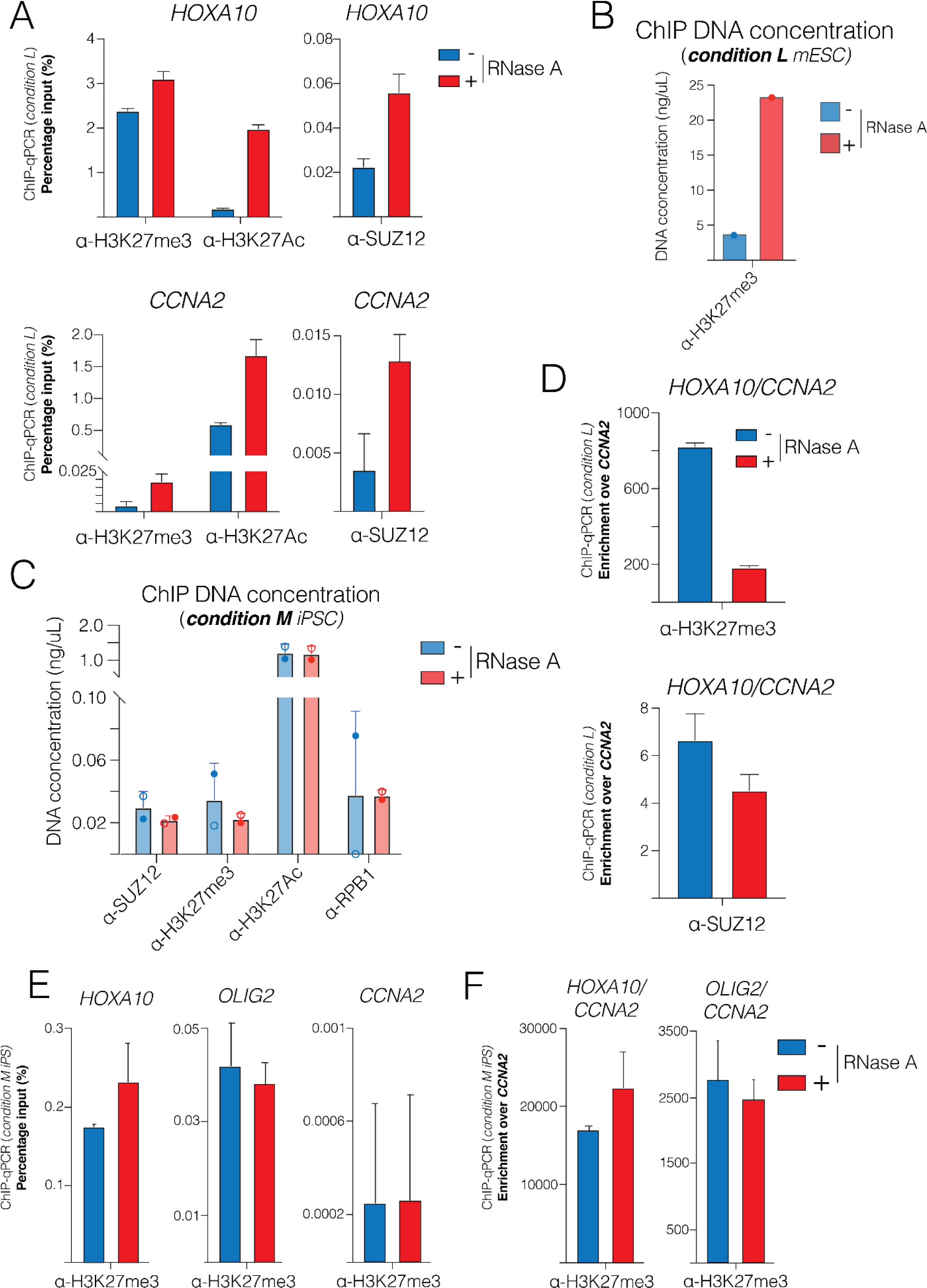
RNA depletion during ChIP leads to a global gain of non-targeted chromatin, which artificially reduces ChIP signals upon data normalisation. (A) ChIP-pPCR analysis of H3K27me3, H3K27Ac and SUZ12 at a representative repressed PRC2 target TSS (*HOXA10*; repressed gene) and active TSS (*CCNA2*; active gene) following RNase A treatment under Condition L in human iPS cells. Enrichment is presented as a percentage of input DNA. Data shown is from an independent replicate to data in Figure 4A, carried out on a different day. (B) The concentration of the DNA immunoprecipitated with an H3K27me3 antibody in the presence and absence of RNase A treatment under condition L in mESC. DNA concentration was measured with a Qubit dsDNA High Sensitivity Assay Kit (ThermoFisher Q32854). (C) The concentration of DNA immunoprecipitated with different antibodies, as indicated, under Condition M in human iPSCs. Qubit readings from two independent rChIP replicates are shown (solid circles: replicate 1; empty circles: replicate 2). (D) ChIP-qPCR analyses from (A) presented as enrichment at *HOXA10* TSS as a percentage of a negative control TSS *CCNA2.* Data shown is from an independent replicate to data in Figure 4C, carried out on a different day. (E) ChIP-qPCR analyses of H3K27me3 following RNase A treatment at representative PRC2 target TSSs (*HOXA10* & *OLIG2)* under condition M in human iPSCs. Enrichment is presented as a percentage of input DNA. (F) ChIP-qPCR analyses of the same samples from (E) presented as enrichment at *HOXA10* or *OLIG2* normalised to a negative control TSS CCNA2.

